# *Fusobacterium nucleatum* induces pancreatic cancer cell proliferation and migration by regulating host autocrine and paracrine signaling

**DOI:** 10.1101/2021.11.19.469245

**Authors:** Barath Udayasuryan, Tam T. D. Nguyen, Ariana Umaña, Raffae N. Ahmad, LaDeidra Monét Roberts, Polina Sobol, Stephen D. Jones, Jennifer M. Munson, Daniel J. Slade, Scott S. Verbridge

## Abstract

Pancreatic ductal adenocarcinoma (PDAC) harbors a complex tumor microbiome that has been implicated in cancer progression and resistance to chemotherapy. Recent clinical investigations uncovered a correlation between high loads of intratumor *Fusobacterium nucleatum* and decreased patient survival. Here we show that healthy and cancerous pancreatic cell lines harboring intracellular *F. nucleatum* secrete increased levels of cancer-associated cytokines including GM-CSF, CXCL1, IL-8, and MIP-3α. We report that GM-CSF (granulocyte-macrophage colony stimulating factor) secretion directly increases the proliferation and migration of pancreatic cancer cells via an autocrine mechanism, notably in the absence of immune cell participation. Furthermore, we show that non-cancerous pancreatic epithelial cells do not exhibit increased proliferation or migration in response to these cytokines, but nevertheless, their secreted cytokines stimulate these responses in cancerous cell lines through paracrine signaling. Our results provide evidence that intratumor *F. nucleatum* in the pancreas elicits an infection-specific cytokine secretion profile from both normal and cancerous cells that adversely contributes to cancer progression through autocrine and paracrine mechanisms. Therefore, these results support the importance of investigating the contributions of both microbiome and host driven processes in pancreatic cancer to guide future therapeutic interventions.

## Introduction

Pancreatic ductal adenocarcinoma (PDAC) is an aggressive malignancy that has a median survival of fewer than 6 months, and a 5-year survival rate of 3-7%^1^. Oncogenic *KRAS* mutation is critical in driving disease progression from early, preinvasive precursor lesions to a malignant and invasive adenocarcinoma^2–4^. Characterized by extensive stromal remodeling and desmoplasia, the PDAC tumor microenvironment (TME) has limited vascularization and poor immune infiltration which makes it highly intractable to chemotherapy^5,6^. Furthermore, the TME is enriched with immunosuppressive cells such as T regulatory (Treg) cells, myeloid-derived suppressor cells (MDSCs), and tumor-associated macrophages (TAMs)^7,8^. This immune suppression is further compounded by tumor-specific microbial species residing within the TME which have been implicated in worse outcomes for patients with PDAC^9–11^.

Recent systematic characterization of intratumoral bacteria in multiple cancer types details a diverse bacterial ecosystem within tumors that is not found in adjacent healthy tissue^12^. These observations have elicited several fundamental questions on the precise role that microbes play within the TME to impact tumorigenesis, cancer progression, and therapy response^13,14^. Little is known on how these bacteria reach the tumor, the survival strategies that they employ within the TME, the host tumor cells’ response to the infection, and if their elimination could augment cancer therapy. Previous studies have revealed positive associations of certain microbes with the development of PDAC. These include *Fusobacterium nucleatum, Porphyromonas gingivalis, Neisseria elongata, Streptococcus mitis*^11,15^, and *Helicobacter pylori*^16^. Moreover, the PDAC tumor microbiome has been shown to influence immune response and can impact chemotherapy^17^. For example, Gammaproteobacteria in PDAC tissue regulates drug resistance to a common chemotherapeutic agent, gemcitabine^18^. However, there is still limited insight into the individual roles these bacteria play in worsening cancer prognosis. Our work aims to shed light on the role of a specific microbe, *F. nucleatum*, and its impact within the PDAC TME.

*Fusobacterium nucleatum* is a Gram-negative, anaerobic, rod-shaped bacterium usually found within the oral cavity. It is a commensal microorganism that plays a supportive role as a bridge-species within biofilms connecting primary and secondary microbial colonizers^19^. However, in certain disease conditions, it can act as an opportunistic pathogen. In addition to participating in inflammatory diseases within the oral cavity, such as in gingival and periodontal disease, it is a risk-factor in preterm birth and intrauterine infections^20^. There is also associative evidence of its role in appendicitis^21^, urinary tract infection, endocarditis^22^, and other respiratory tract infections^23^. Moreover, *F. nucleatum* is increasingly recognized as an oncomicrobe due to its disproportionate presence in several cancers and association with a worse prognosis. These cancers include colorectal cancer^24,25^, esophageal cancer^26^, pancreatic cancer^27,28^, and Lauren’s diffuse type gastric cancer^29^.

To date, most mechanistic studies of *F. nucleatum* have focused on its role in colorectal cancer (CRC). *F. nucleatum* uses its surface adhesin, Fap2, to initially bind to host Gal/Gal-NAc^30^ sugar residues found abundantly in CRC, which then drives entry into host cells and intracellular colonization. In fact, Gal/Gal-NAc is highly expressed in a wide variety of tumors and may act as a key factor in bacterial homing to these sites^31–33^. In addition, the adhesin FadA has been shown to bind to E-cadherin, which activates β-catenin and stimulates downstream signaling that promotes carcinogenesis^34^. However, there is still a critical knowledge gap in our understanding of how *F. nucleatum* adapts to the TME or continues to influence cancer phenotypes once colonizing the TME, to exacerbate the aggressiveness of both early and late-stage cancers^35^. Moreover, it has been shown that *F. nucleatum* may travel within the primary cells to distant tumor metastatic sites^36^ and could contribute to pre-metastatic niche formation. We have previously shown that *F. nucleatum* infects host cells in a Fap2-driven mechanism, and leads to the secretion of potent cytokines, IL-8 and CXCL1, which we demonstrated to directly increase cellular migration of a colorectal cancer cell line, HCT116^37^, which may directly facilitate metastatic spread of CRC.

Emerging evidence has tied *F. nucleatum* to increased mortality from pancreatic cancer. Mitsuhashi et al. initially uncovered the association of *Fusobacterium* species in PDAC^28^. They tested cancer tissue specimens from 283 patients with PDAC and identified an 8.8% detection rate of *Fusobacterium* species and observed highly significant mortality in the *Fusobacterium* positive group. Gaiser et al. identified an enrichment of oral bacterial taxa including *F. nucleatum* and *Granulicatella adiacens* in cyst fluid from intraductal papillary mucinous neoplasms (IPMNs) with high-grade dysplasia (HGD)^27^. Additionally, Alkharaan et al. identified circulating IgG reactivity to *F. nucleatum* and high salivary IgA reactivity to *F. nucleatum*, and Fap2 of *F. nucleatum* in patients with HGD^38^. Further corroboration comes from the study by Nejman et al. which identified *F. nucleatum* in pancreatic cancer with a prevalence of 15-20% from the 67 tumor samples that were tested^12^. These observations suggest a disproportionate presence of *F. nucleatum* within the pancreatic TME, warranting further research into its function in pancreatic cancer pathogenesis.

A distinguishing feature of PDAC is that tumor cells form only a small fraction of the TME. In fact, studies have shown how PDAC progression and resistance to therapy is considerably driven by its dense stroma, which can constitute up to 80% of the tumor^39^. It consists of multiple non-oncogenic cell types including cancer associated fibroblasts, pancreatic stellate cells, immune cells, and endothelial cells that shape the behavior of PDAC cells, by remodeling the extracellular matrix, promoting epithelial-mesenchymal transition (EMT), and fostering aberrant cell signaling^40,41^. The dense fibrosis and limited vascularization hence formed creates a stiff and hypoxic TME that could provide a favorable niche for anaerobic organisms such as *F. nucleatum*.

Our previous findings from *F. nucleatum* infection of CRC cells showed that bacterial infection can exert local effects on cancer cells without the involvement of immune cells^37^. We hypothesized that *F. nucleatum* may play an analogous role in PDAC to impact tumor progression. There have been few if any studies to our knowledge that have investigated the phenotype of *F. nucleatum* infected healthy and cancerous pancreatic cell lines, and it is unclear if the infection of these cells is Fap2-driven as in the case of CRC. We therefore sought to investigate the specific role of *F. nucleatum* in pancreatic cancer and identify key secretory factors from host cells that could contribute to worse outcomes in patients harboring this bacterium in their tumors.

## Results

We carried out our studies using four pancreatic cancer cell lines and a primary normal pancreatic epithelial cell line, the details of which are shown in **Supplementary Table 1**^42^. These lines were selected as they were extensively characterized, and exhibit differences in sex, metastasis, and differentiation, which could impact their response to *Fusobacterium* infection. The data from BxPC3 and Panc1 cell lines are shown in the main text, and the data from HPAC and Capan1 cell lines can be found in the Supplementary Material.

### Fusobacterium binds and invades BxPC3 and Panc1 cells through a Fap2-driven mechanism

We first confirmed that *F. nucleatum* can bind and invade pancreatic cancer cells. Scanning electron microscopy (SEM) imaging enabled direct visualization of the various modes of binding and invasion of *F. nucleatum subsp. nucleatum* ATCC 23726 (which will be subsequently referred to as *Fnn* in this article) into BxPC3 cells (**Figure 1A**). We further confirmed the intracellular localization of *Fnn* within BxPC3 cells through confocal imaging and flow cytometry (**Figure 1B**). We sorted the cells based on the intensity of fluorescence observed and imaged them in high resolution z-stacks to visualize the number of intracellular bacteria within each cell. Based on our work and previous research highlighting the importance of Fap2 in *F. nucleatum* binding, we hypothesized an analogous interaction with pancreatic cancer cells. Using flow cytometry, we identified that the Fap2 surface adhesin deletion mutant (*Fnn Δfap2*), previously created and used in our work^37,43^, infects both BxPC3 and Panc1 cells at a significantly decreased rate compared to the wild-type *Fnn* (**Figure 1C**).

**Figure 1.**
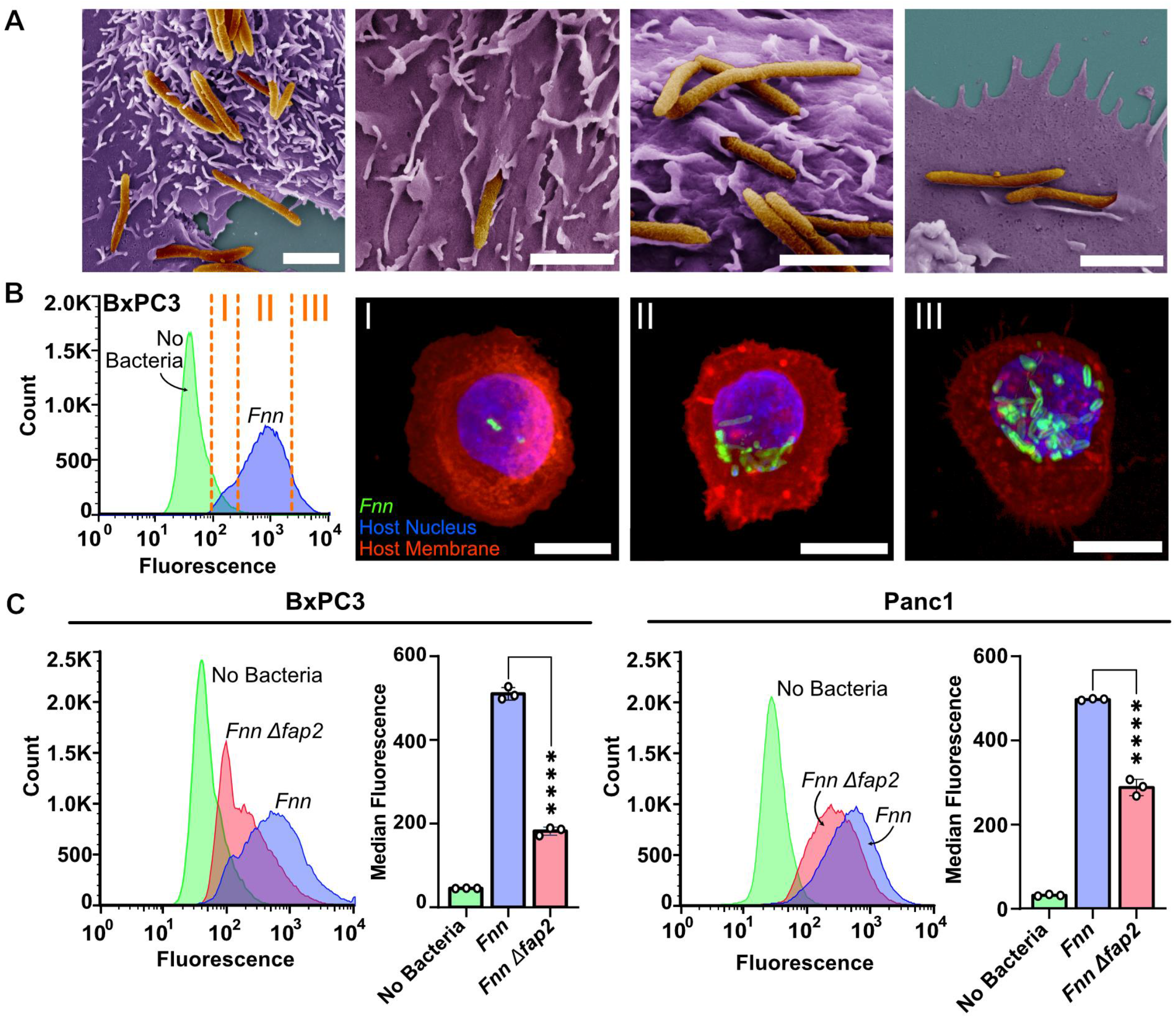
*Fusobacterium nucleatum* binds to and invades pancreatic cell lines. **A.** Scanning electron microscopy images of *F. nucleatum subsp. nucleatum 23726* (*Fnn*) invading BxPC3 pancreatic cancer cells line through multiple points of adherence to the cell membrane. Pseudocolored with: bacteria in orange, host cells in purple (scale bar: 2μm). **B.** Flow cytometry analysis of BxPC3 cells without bacteria (No Bacteria) and with infection with *Fnn* at 50:1 MOI demonstrating an increase in fluorescence of infected cells. The cells were further sorted and collected based on three levels of fluorescence intensity (I,II,III), corresponding to increased intracellular bacterial loads. These cells were plated and imaged on a confocal microscope to identify relative levels of infection. Blue: host cell nuclei stained with DAPI; green: *Fnn* membrane stained with FM 1-43 FX; red: host cell membrane stained with MemBrite 568 (Biotium) (scale bar: 10 μm). **C.** Flow cytometry analysis of *Fnn* vs *Fnn Δfap2* infection of BxPC3 and Panc1 cells at 1 hour after a 50:1 MOI shows a significantly slower infection of *Fnn Δfap2* indicating Fap2 driven intracellular invasion. N = 3 independent experiments, compared using ordinary one-way ANOVA, followed by Dunnett’s multiple comparisons test. P-value significance denoted by ns (not significant) for P>0.05, and **** for P≤0.0001.

### Host pancreatic cancer cells secrete specific cytokines upon *Fusobacterium* infection

Once we confirmed host cell binding to *Fnn* and subsequent intracellular invasion into host cells, we then analyzed the host cells’ response to infection. Western blot arrays were used to identify infection-induced cytokine secretions from the pancreatic cancer cell lines tested in four conditions: (a) No bacteria (b) infection with *Escherichia coli*, (c) infection with *Fnn* and (d) infection with *Fnn Δfap2*. Heat maps were constructed based on the raw intensities (**Supplementary Figure 1**) and the percent fold change in protein secretion compared to the No Bacteria condition was determined (**Figure 2A**). While pancreatic cancer cell lines are highly secretory without infection, we identified *Fnn* infection specific increases in secretion for GM-CSF, CXCL1, IL-8, and MIP-3α. There was minimal or no change in secretion in response to infection by *E. coli*. ELISAs were performed to confirm and quantify the secretion of these cytokines, and significant increases were noted for the four cytokines when compared to the No Bacteria condition for BxPC3 and Panc1 cells (**Figure 2B**, **Supplementary Figure 2**). There was a significant increase in IL-8 and CXCL1 secretion upon infection with *E. coli* for BxPC3 cells but not for Panc1 cells. Furthermore, there was a significant increase in the secretion of GM-CSF in BxPC3 cells upon infection with *E. coli* with respect to No Bacteria, but was significantly lower (p≤0.001) compared to GM-CSF secretion elicited by infection with *Fnn* and *Fnn* Δ*fap2*. Notably, there was minimal change in response to measured cytokine values between infection with *Fnn* and *Fnn* Δ*fap2*.

**Figure 2:**
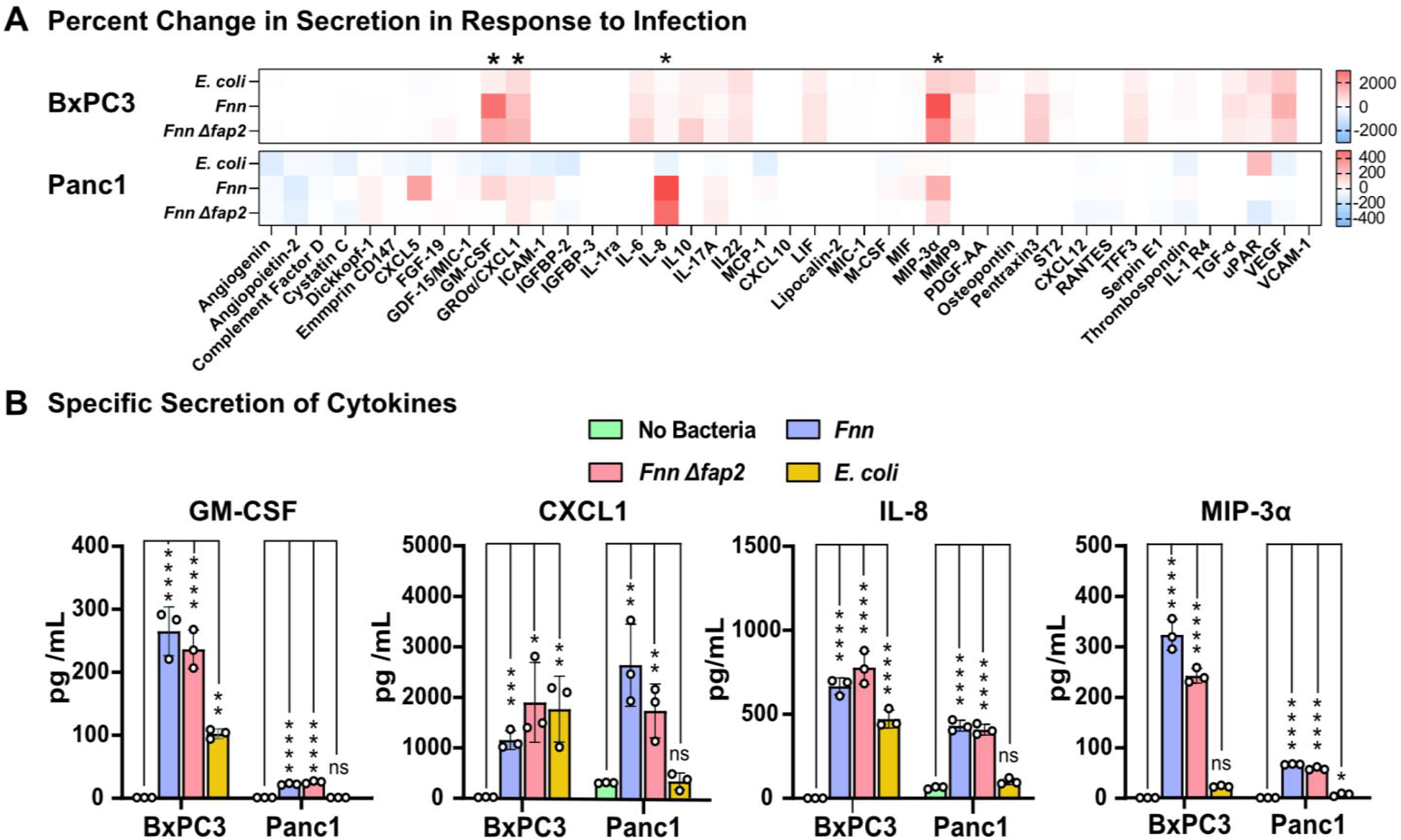
*Fnn* infection elicits secretion of specific cytokines from host cells. **A.** Heat maps depicting the percent change in cytokine secretion in response to infection in comparison to the non-infected samples of BxPC3 and Panc1 cell lines tested using the Human XL cytokine array (R&D Systems Inc.) upon infection with *E.coli, Fnn*, and *Fnn Δfap2*. Using these results, we identified the specific increase in secretion of GM-CSF, CXCL1, IL-8, and MIP-3α (marked with an asterisk). **B.** Measured values of selected cytokines using ELISA confirming specificity to *Fnn* infection. N = 3 independent experiments, compared using ordinary one-way ANOVA, followed by Dunnett’s multiple comparisons test. p-value significance denoted by ns (not significant) for P>0.05, * for P≤0.05, ** for P≤0.01, *** for P≤0.001, and **** for P≤0.0001.

### GM-CSF increases the proliferation of pancreatic cancer cells

We hypothesized that the four cytokines we identified from the cytokine arrays could be involved in impacting the proliferation of pancreatic cells; a common feature in aggressive cancers. We first observed their impact on proliferation individually on BxPC3 cells using an XTT assay (**Supplementary Figure 3**). From this preliminary assay, we discovered that GM-CSF at a concentration of 200 pg/mL increased the proliferation of BxPC3 cells, while the other cytokines, IL-8, CXCL1, and MIP-3α, did not. In both BxPC3 and Panc1 cells, we observed increased proliferation of cells when infected with *Fnn*. Furthermore, conditioned media obtained from infected cells after a 4-hour infection was sufficient to induce the proliferation of uninfected cells. This was confirmed by two independent assays to quantify proliferation (BrdU and XTT, **Figure 3A, 3B**). The addition of recombinant human GM-CSF (rhGM-CSF) at 200 pg/mL was sufficient to increase the proliferation of BxPC3 and Panc1 cells (**Figure 3C**). To confirm that GM-CSF was primarily responsible for the proliferative effect, we depleted GM-CSF from the conditioned media using biotinylated anti-GM-CSF antibodies to capture free GM-CSF and used streptavidin-coated magnetic particles to capture and spin down the cytokines and observed a concomitant decrease in proliferation (**Figure 3D**). Since Panc1 cells showed lower but measurable secretion of GM-CSF, we supplemented 200 pg/mL rhGM-CSF to these cells, and depleted rhGM-CSF from the media once again and confirmed that GM-CSF directly contributed to an increase in cell proliferation of this pancreatic cancer cell line (**Figure 3E**). This last assay indicates that exogenous GM-CSF within the TME can directly impact the proliferation of pancreatic cancer cells.

**Figure 3:**
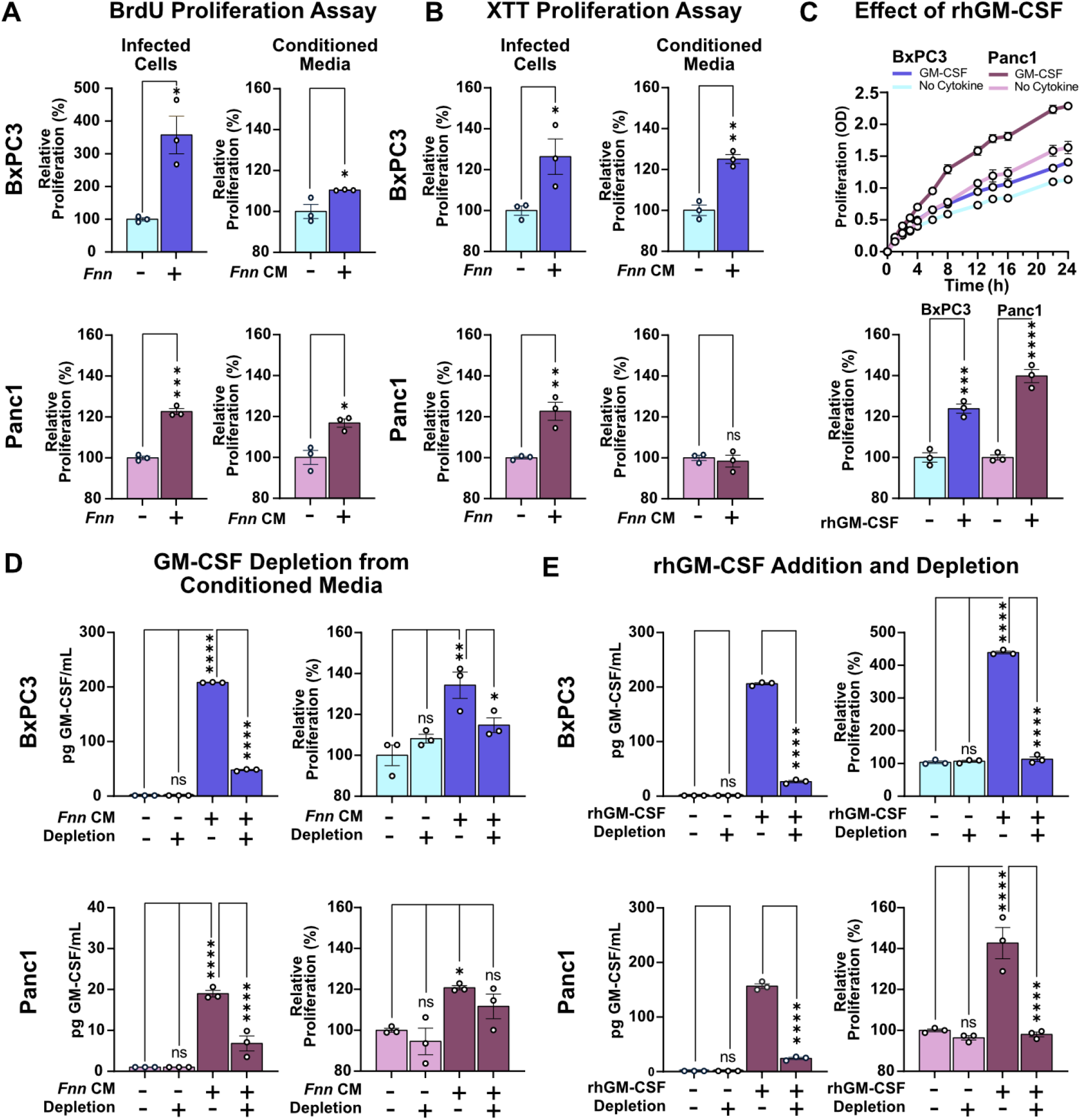
GM-CSF induces pancreatic cell proliferation. **A, B.** BrdU and XTT assays both independently confirmed that *Fnn* infected host BxPC3 and Panc1 cells have increased proliferation compared to non-infected cells. This increase in proliferation of non-infected cells is sustained in the presence of conditioned medium (*Fnn CM*) isolated from infected cells. **C.** Addition of recombinant human GM-CSF (R&D Systems 200 pg/mL) to BxPC3 and Panc1 cells significantly increases their proliferation. **D.** GM-CSF depletion from conditioned media obtained from *Fnn* infected cells significantly reduced the relative proliferation of BxPC3 and Panc1 cells. **E.** GM-CSF depletion after supplementing cells with exogenous rhGM-CSF also reduces cell proliferation of BxPC3 and Panc1 cells. N = 3 independent experiments, compared using student’s t-test, ordinary one-way ANOVA, followed by Dunnett’s multiple comparisons test. p-value significance denoted by ns (not significant) for P>0.05, * for P≤0.05, ** for P≤0.01, *** for P≤0.001, and **** for P≤0.0001.

### Infected Cells and non-infected cells exhibit increased migration in response to *Fnn* induced host cell secretion

Since several studies have implicated the four upregulated cytokines we identified in cell migration, we hypothesized that the *Fnn* induced host secretion would impact pancreatic cancer cell migration. We determined the migration of *Fnn* infected and uninfected host cells in an in vitro Transwell co-culture model. BxPC3 cells were infected with *Fnn* at 50:1 MOI for 4 hours after which they were collected and added to the top chamber of a Transwell with a membrane that consisted of 8μm pores. *Fnn* infected BxPC3 cells trans-migrated across the membrane in a Transwell assay at a significantly increased rate compared to uninfected cells over 16 hours (**Figure 4A**). In another assay, we collected conditioned media from the 4-hour infection of host cells with *Fnn* at 50:1 MOI, filtered and concentrated it 16x, then added the concentrated media to the outer (lower) chamber of the transwell. We observed significantly greater migration of uninfected BxPC3 cells from the top of the membrane to the bottom in response to concentrated conditioned media obtained from infected cells than in response to concentrated conditioned media obtained from uninfected cells over 16 hours (**Figure 4B**). The short time period of 16 hours and using 1% serum supplemented media restricted the contribution of cellular proliferation in this assay.

**Figure 4:**
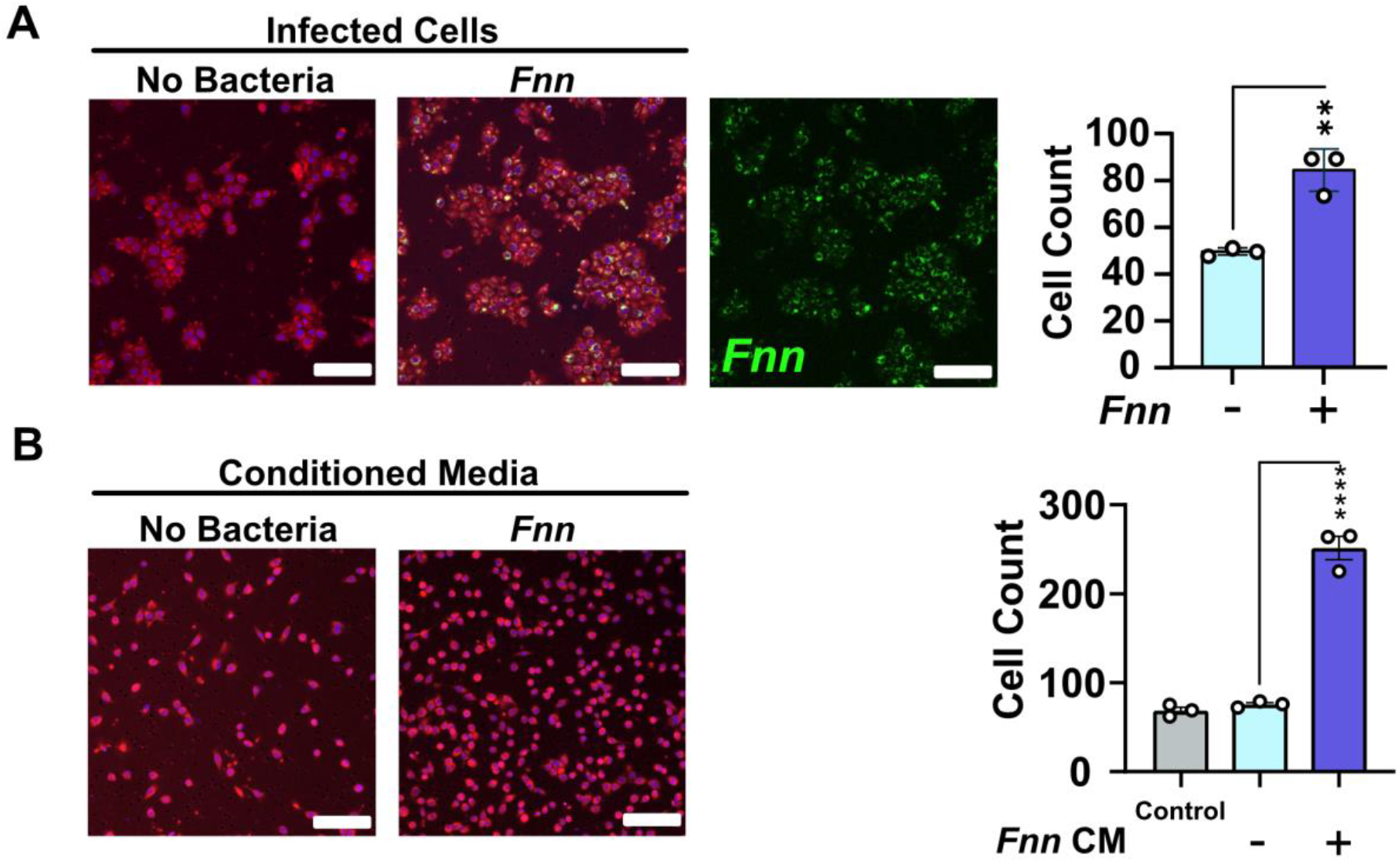
Increased cell migration observed for both *F. nucleatum* infected cells and non-infected cells in response to conditioned media obtained from *F. nucleatum* infected cells. Representative images of Transwells with BxPC3 cells that have migrated over 16 hours in response to the control with 0.5% FBS, and conditioned media obtained from cells not infected and infected with *Fnn*. (Staining: Red: Cell Tracker Red, Blue: DAPI, Green: *Fnn* with FM 1-43X) **A.** BxPC3 cells infected with *Fnn* show significantly increased migration compared to uninfected cells. **B.** BxPC3 cells show significantly increased migration in response to conditioned media obtained from infected cells (Staining: Red: Cell Tracker Red, Blue: DAPI) (scale bar = 100μm). N = 3 independent experiments, compared using ordinary one-way ANOVA, followed by Dunnett’s multiple comparisons test. P-value significance denoted by ns (not significant) for P>0.05, * for P≤0.05, ** for P≤0.01, *** for P≤0.001, and *\*\*\*\** for P≤0.0001.

### Normal pancreatic epithelial cells secrete cytokines upon *F. nucleatum* invasion that stimulate proliferation and migration of cancer cells

We next studied the infection of a normal pancreatic epithelial cell line (which will be referred to as nPEC in this text) and compared its response with that from infection of pancreatic cancer cell lines. Flow cytometry analysis of nPECs upon infection with *Fnn* and *Fnn Δfap2* indicated that the infection occurs at the same rate for both strains (**Figure 5A, Supplementary Figure 4**). Next, ELISA of conditioned media from nPECs confirmed a significant secretion of GM-CSF, CXCL1, IL-8, and MIP-3α in response to *Fnn* and *Fnn Δfap2* infection, which was absent upon infection with *E. coli* (**Figure 5B**). There was also reduced secretion of GM-CSF upon infection with *Fnn Δfap2* when compared to infection with *Fnn*. Proliferation assays were subsequently performed and in contrast to the cancer cell lines, nPECs did not increase proliferation upon infection or in the presence of conditioned media obtained from infected nPECs (**Figure 5C**). Similarly, the nPECs did not proliferate in the presence of 200pg/mL recombinant hGM-CSF (**Figure 5D**). However, when conditioned media obtained from infected nPECs was added to BxPC3 cells, it stimulated BxPC3 cells to proliferate (**Figure 5E**). Next, we studied migratory response in a Transwell assay and observed that infection of nPECs did not impact migration in comparison to uninfected nPECs (**Figure 5F**). Furthermore, we observed that conditioned and concentrated media from infected nPEC cells did not impact nPEC migration, but significantly increased the cell migration of BxPC3 cells (**Figure 5G**).

**Figure 5:**
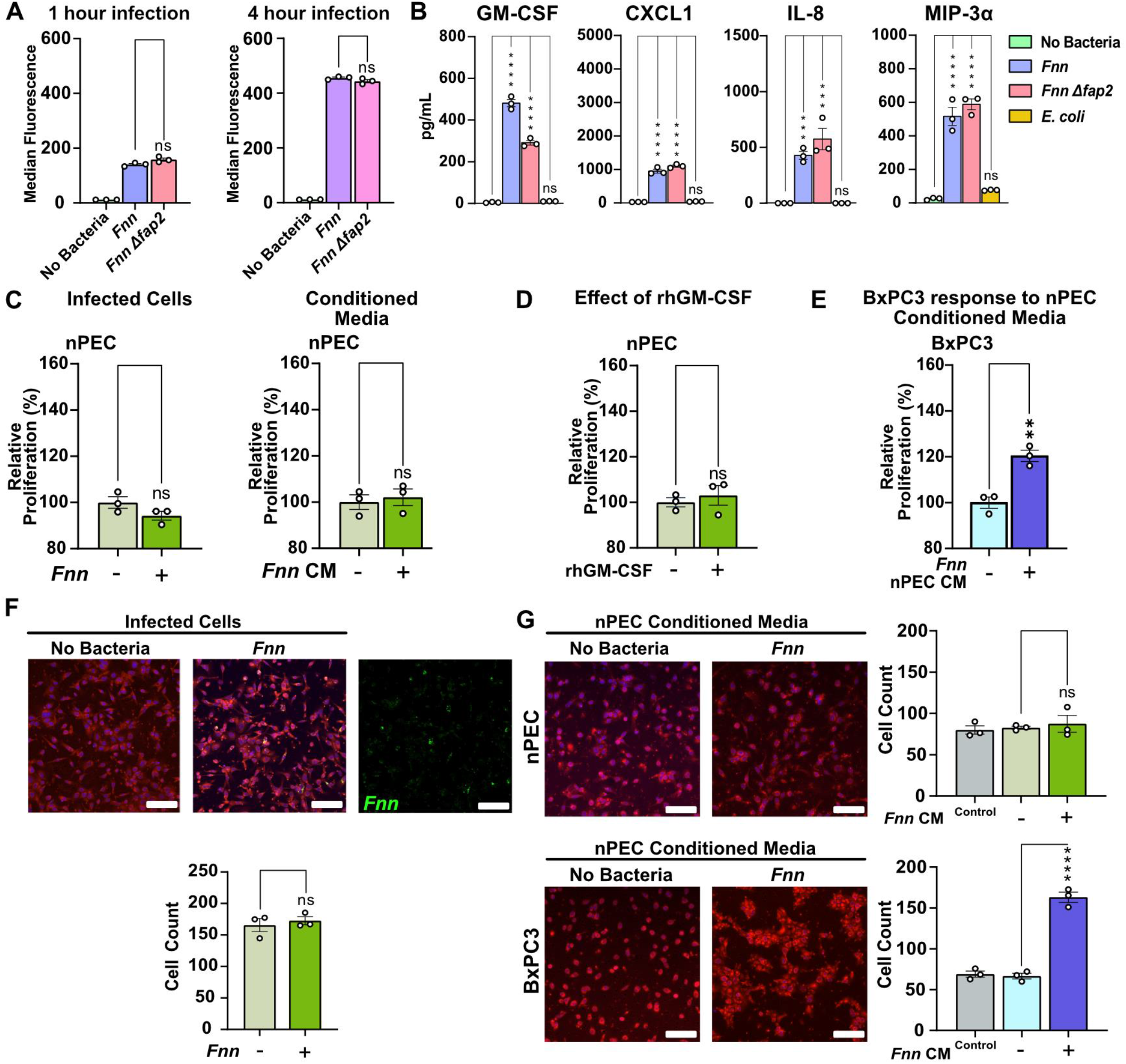
Normal Pancreatic Cells response to *F. nucleatum* infection. **A.** Flow cytometry analysis of nPEC infection with *Fnn* and *Fnn Δfap2* at 1 and 4 hours indicates equal rate of infection. **B.** Increased secretion of GM-CSF, CXCL1, IL-8, and MIP-3α, from nPECs upon infection with *Fnn, Fnn Δfap2*, and *E. coli* compared with the uninfected condition. **C.** XTT proliferation assays show no increase in the proliferation of infected nPECs, nPECs in response to conditioned media obtained from infected cells **D.** nPECs do not increase their proliferation in response to rhGM-CSF. **E.** BxPC3 cells show increased proliferation in response to conditioned media obtained from infected nPECs. Transwell migration assay shows that infected nPECs do not increase migration in response to infection. **F.** Infected nPEC do not show an increase in migration **G.** Transwell migration assays with conditioned and concentrated media obtained from infected nPEC cells do not stimulate migration of nPEC cells but stimulate BxPC3 cells to migrate at a significantly increased rate (scale bar: 100 μm). N = 3 independent experiments, compared using ordinary one-way ANOVA, followed by Dunnett’s multiple comparisons test. P-value significance denoted by ns (not significant) for P>0.05, * for P≤0.05, ** for P≤0.01, *** for P≤0.001, and **** for P≤0.0001.

## Discussion

Motivated by our prior work on the role of *F. nucleatum* in inducing cytokine secretion and cellular migration in the CRC TME, we hypothesized that *F. nucleatum* may play analogous critical roles within the PDAC TME, potentially directly acting to enhance cancer progression. We sought in this study to leverage in vitro co-culture models to begin to define and quantify these roles that would be challenging to identify and dissect in vivo. We confirmed the intracellular invasion of this bacteria in pancreatic normal and cancer cell lines and have identified the specific secretion of cytokines that may adversely affect tumor progression, by increasing tumor cell proliferation and enhancing tumor cell migration.

Using flow cytometry, SEM, and confocal imaging, we first confirmed intracellular invasion and localization of *F. nucleatum* in pancreatic cell lines. *F. nucleatum* is well-known to bind and invade a variety of epithelial cell types,^19,44^ although in most cases, exact mechanisms for organ specific cell binding and invasion need to be investigated further. Among the host of binding adhesins expressed on *F. nucleatum*, a critical adhesin and lectin involved in binding is the 390 kDa outer membrane protein Fap2. Fap2 is a member of the type 5a secreted autotransporter family of proteins which are critical virulence factors expressed by Gram-negative bacteria. This protein functions in adhesion, cell-cell aggregation, biofilm formation, and bacterial invasion. In CRC, it has been shown that Fap2 binding to human inhibitory receptor TIGIT on natural killer (NK) cells and the tumor itself can protect tumors from being targeted by NK cells and also inhibits T cell function by causing tumor-immune evasion^45,46^. *F. nucleatum* binding to tumor cells is also mediated by Fap2, evidenced in CRC30 and breast cancer^32^. Its target is Gal-GalNAc (D-galactose-β(1-3)-N-acetyl-D-galactosamine) that is expressed at high levels in adenocarcinomas, including pancreatic cancer^31,33^. Furthermore, it was identified that the Fap2 lectin domain can be inhibited by galactose and galactose derivative molecules^47,48^. Our results reveal significantly decreased infection of BxPC3 and Panc1 cells by the Fap2 deficient *F. nucleatum* in comparison to the wild-type, indicating that Fap2 binding plays a critical role in the intracellular invasion of the bacterium in pancreatic cancer as well. Notably, this difference is not observed upon binding and invasion of normal pancreatic cells by the *fap2* deletion mutant, indicating that healthy pancreatic cell invasion is not driven by Gal-GalNAc docking, and perhaps adhesins other than Fap2 contribute to binding normal epithelial cells through undiscovered protein-protein interactions. A frequently proposed approach to inhibit *F. nucleatum* binding to tumor cells has been to use galactose-containing compounds to inhibit Fap2 lectin domain interaction. However, if normal tissue adjacent to the tumor can also be infected at the same rate, targeting Fap2 interactions would not be effective in stopping healthy tissue secretion of pro-cancerous cytokine profiles that contribute to proliferation and migration through paracrine mechanisms. Therefore, we acknowledge that further research is required to identify and characterize additional adhesins *F. nucleatum* uses to bind to and invade normal cells to block these cross-talking pathways in cancer.

The next focus of our study was the pancreatic cancer cells’ response to *F. nucleatum* infection, specifically host cell secretions of cytokines. Cytokines are soluble secreted proteins usually with a molecular weight <30 kDa that mediate cell-cell communication. They bind to high-affinity receptors present on cell membranes that activate paracrine or autocrine signaling pathways to stimulate host cells in response to injury, cellular stress, inflammation, and immune cell recruitment. Cancer cells have hijacked these signaling mechanisms and exploit them to promote angiogenesis, metastasis, differentiation, immune suppression, and drug resistance. Thus, there is considerable focus on developing cancer immunotherapies utilizing or targeting cytokines^49^. However, there is immense difficulty in delineating the individual roles of cytokines due to their synergistic and antagonistic mechanism of action, which additionally have complex temporal and concentration dependence. Our findings from the cytokine arrays identified *F. nucleatum* specifically induced the secretion of GM-CSF, CXCL1, IL-8, and MIP-3α by BxPC3, Panc1, and HPAC cells, and increased IL-6, GM-CSF, and MIP-3α secretion in addition to the constitutive secretion of IL-8 and CXCL1 by Capan1 cells. Pancreatic cancer cells are highly secretory, and in addition to other constitutively secreted factors, these cytokines are well-known players in PDAC tumor progression and immune suppression.

The cytokines that we have identified in this study in response to *F. nucleatum* infection have been implicated in various forms of cancer^50^. Notably, IL-8 is known to regulate multiple aspects of tumor progression^51^, CXCL1 has been implicated in chemoresistance and metastasis^52^ as well as pre-metastatic niche formation in CRC^53^, and MIP-3α (or CCL20) has multifaceted roles within the TME^54^. MIP-3α production has also been observed in *F. nucleatum* infection of human oral epithelial cells^55^. Intriguingly, GM-CSF is known to exhibit both stimulatory and suppressive effects on tumor progression^56,57^ and has been shown to play a role in liver metastasis^58^. These observations still do not reveal if this is an active or passive process (referring to increased expression versus cytokine release from cells), but the selectivity of secretion indicates the activation of specific pathways within the host cells. The ubiquitous presence of these cytokines in several cancer types suggests their potential role as cancer biomarkers^59^.

Previous studies have also revealed roles for these cytokines in pancreatic cancer. It has been shown that KRAS-induced upregulation of GM-CSF can promote the development of pancreatic neoplasia^60^. GM-CSF simultaneously plays a role in oncogenic activation and the evasion of antitumor immunity in PDAC. Moreover, it is an important regulator of inflammation and immunosuppression within the TME, such as its characterized role in the recruitment of immunosuppressive Gr1^+^CD11b^+^ myeloid cells^61^. GM-CSF secreted by mesenchymal stem cells within the PDAC stroma has been previously shown to promote tumor cell growth and metastasis^62^. These multiple roles have been tied to worsening patient outcomes. In fact, in a study of 68 patients with PDAC, GM-CSF high populations had shown a significantly lower survival rate (a median survival of 13.9 months compared to 25.7 months for GM-CSF low population)^63^. MIP-3α is known to promote Panc-1 cell invasion in vitro by binding to its CCR6 receptor, complemented with the up-regulation of matrix metalloproteinases (MMP9)^64^. Furthermore, Takamori et. al have shown that IL-8 and CXCL1 autocrine signaling increases the proliferation of Capan1 cells^65^. These studies further corroborate our results that *F. nucleatum* induced host cell secretion of these cytokines contribute adversely to cancer progression.

Our results reveal an additional role for GM-CSF in pancreatic cancer. We studied the individual and collective impact of the identified cytokines using in vitro co-culture assays of proliferation and migration which are common features of aggressive cancers. We demonstrated that cytokines secreted from infected cancer cells can directly impact other uninfected cancer cells. We first noted increased proliferation of infected cells. To ascertain that host cell cytokine secretions could play a role to explain this observation, we isolated conditioned media from the *F. nucleatum* infected cancer cells and found that it was sufficient to increase proliferation of the pancreatic cancer cells, and this effect was partially negated when we depleted GM-CSF from the media. We also demonstrated that the addition of exogenous recombinant GM-CSF increased cell proliferation and its depletion negated the observed effect. Together, these observations define an independent role of GM-CSF in BxPC3 and Panc1 cells, to directly contribute to cancer progression by impacting proliferation.

We additionally observed increased cell migration of infected cells compared to uninfected cells, and hypothesized that secreted cytokines could account for this altered phenotype. We demonstrated that conditioned and concentrated media obtained from *F. nucleatum* infected cells was sufficient to significantly increase the cellular migration of uninfected cells. It is essential to note that this conditioned media contained a mix of other secreted cytokines such as IL-6, IL-10, TNFα, and VEGF, albeit in lower concentrations, in addition to high levels of GM-CSF, CXCL1, IL-8, and MIP-3α, as revealed from the cytokine arrays. Nevertheless, the resulting cell migratory response arising from a synergistic fusion of these cytokine interactions was specific to *F. nucleatum* infection. These observations indicate that infected cancer cells can impact both infected as well as uninfected cancer cells within the PDAC TME through both autocrine and paracrine signaling, an important potential mechanism whereby the impact of relatively low numbers of bacteria in vivo^66^ may be amplified locally through host cell crosstalk.

While the influence of stromal cells such as tumor-associated fibroblasts and pancreatic stellate cells are well characterized in PDAC^40,67–70^, the role of normal epithelial cells has not received as much scrutiny. Our results show that they too can regulate the fate of tumor cells. We show *F. nucleatum* infects normal pancreatic epithelial cells using a Fap2-independent mechanism, but that this infection still results in the secretion of the same cytokines released by infected PDAC cells. These cytokines, which do not elicit a proliferative response from the normal cells, are able to impact tumor proliferation and migration. Cytokines such as IL-8 and CXCL1 are well known players in formation of pre-metastatic niches^53,71^ priming organs and tissues to attract circulating tumor cells (CTC), break dormancy, and establish micrometastatic sites. Furthermore, as tumor cells increase migration in response to cytokines, infected normal cells can stimulate the expansion of the tumor, providing increased opportunity for the tumor cells to access vasculature and disseminate. This cell migration can be either individual or collective in multicellular clusters of cells as we have observed with BxPC3 cells. Collective cell migration has been considered far more detrimental due to tumor cells enhanced pro-survival adaptations and increased metastatic potential^72,73^. In fact, many of the mechanisms that impact metastatic progression of the cancer also contribute to enhanced therapeutic resistance^74^. Thus, the crosstalk between normal and tumor cells through both paracrine and autocrine signaling could lead to the evolution of potentially enhanced tumor subpopulations with increased mobility and can contribute to worse prognosis. Moreover, infection of normal cells also suggests that *F. nucleatum* infections in the vicinity of the tumor can extend to adjacent tissues and organs.

There have been hints as to the mechanisms by which *F. nucleatum* can impact cancer cell proliferation. However, these specific interactions have not been studied in pancreatic cancer. In CRC, a proposed mechanism of action is that FadA adhesin protein from *F. nucleatum*, upon binding to E-cadherin, enables host cell invasion that promotes E-cadherin phosphorylation resulting in increased β-catenin translocation to the nucleus and subsequently increased transcription of Wnt signaling genes^34^. Furthermore, FadA upregulates the expression of annexin A1 (ANXA1), a phospholipid-binding protein, in CRC cells by binding E-cadherin, and ANXA1 can engage β-catenin to activate cyclin D1 to promote cellular proliferation^75^. In mice, it was found that *F. nucleatum* increases CRC proliferation by activating TLR4 signaling, NFκB activation, and the upregulation of miR21^76^. Whether similar pathways are activated in pancreatic cancer remains to be determined. Interestingly, BxPC3 cells do not contain the driver KRAS mutation found in over 90% of pancreatic cancer cells, which is a critical nexus for multiple intracellular signaling pathways related to cell proliferation and differentiation. However, BxPC3 cells still contain chronically active effector pathways related to BRAF^4^ that are further tied to ERK and MAPK pathways that impact proliferation^77^. In fact, GM-CSF is known to activate MAPK and NFκB signaling in other cell types^56,78,79^. In CRC, it was identified that patients with mesenchymal tumors with Fusobacteria have a worse prognosis than patients with a more epithelial subtype^35^. Future work should therefore also investigate *F. nucleatum* impact based on the context of the genomic constitution or molecular subtype of pancreatic cancer^80–82^.

**Figure 6:**
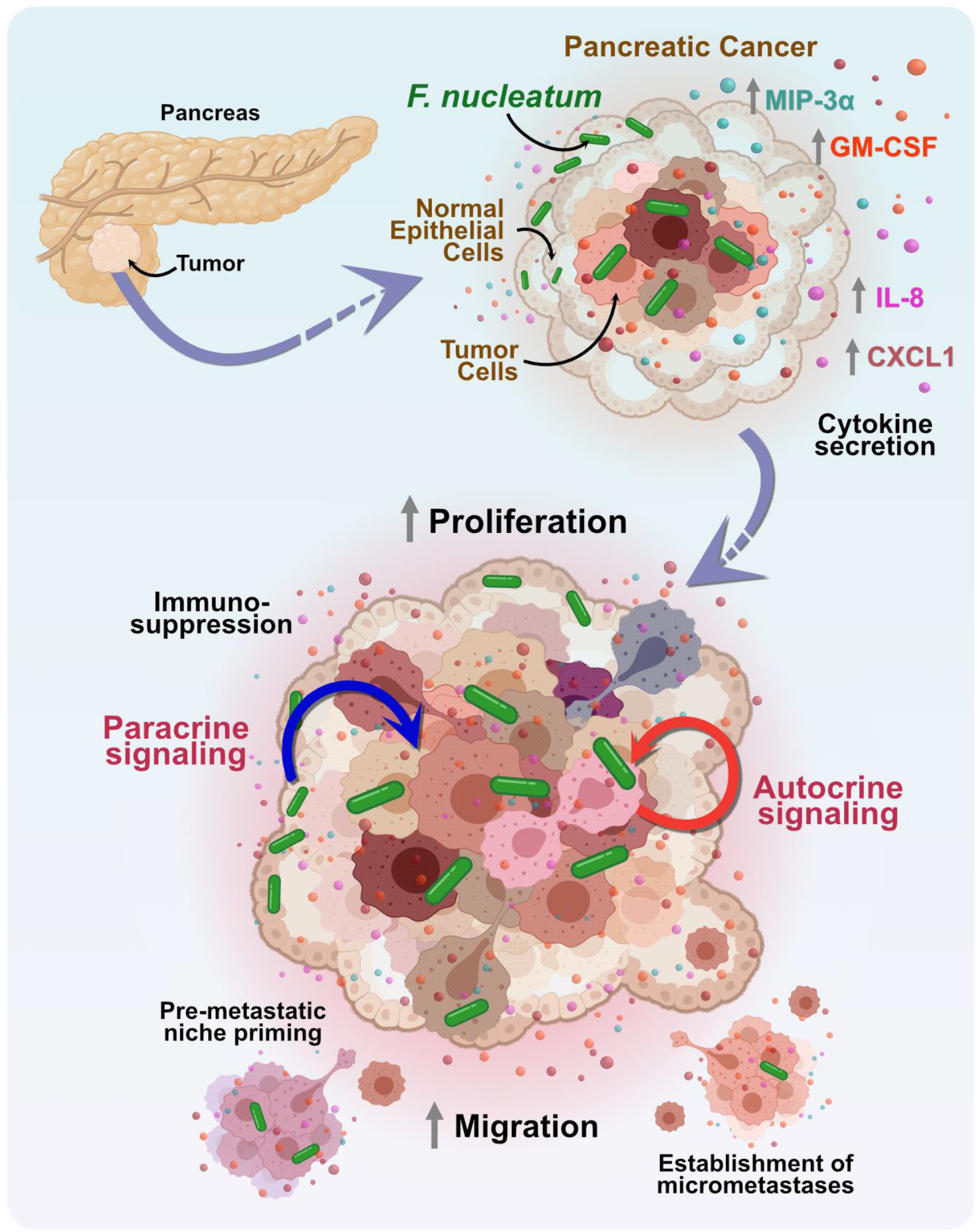
A model of pancreatic cancer response to *F. nucleatum* infection induced cytokine signaling. *F. nucleatum* binds to pancreatic cancer cells in a Fap2-driven mechanism and induces the specific secretion of cytokines, GM-CSF, CXCL1, IL-8, and MIP-3α. These cytokines play a role to enhance proliferation and migration of pancreatic cancer cells through paracrine and autocrine signaling. In addition to known mechanisms of immunosuppression, these responses together adversely impact cancer progression by stimulating metastatic spread, pre-metastatic niche priming, and the establishment of micrometastases.

Tissue-resident microbes can influence cancer susceptibility in diverse and complex ways, for example by modulating inflammation and immune response, altering the metabolic profiles within tumors, and promoting DNA damage. In addition, we now show that host cell secretions upon infection can confer altered proliferative and migratory phenotypes to tumor cells to further exacerbate the cancer. A planned limitation of this study is the lack of response of immune cells to increased cytokine secretion. However, we did this to show that autocrine and paracrine signaling from the tissue cells themselves is sufficient to exacerbate tumor growth. Our work is restricted to the single bacterium *F. nucleatum*, but we note that this bacterium could synergistically act in coordination with other microbes identified within the pancreatic TME.

While PDAC remains one of the most therapy-resistant cancer types, treatments that consider the broader microenvironment of PDAC may provide a more holistic approach, and provide more beneficial outcomes for patients. Our work reveals for the first time that the PDAC-resident bacterium *F. nucleatum* plays a role in directly enhancing tumor cell proliferation and migration, two key hallmarks of PDAC as well as other cancers. The infection of normal cells surrounding the TME further extends the impact of *F. nucleatum* to surrounding tissues and organs, demonstrating that although *F. nucleatum* may not directly transform normal cells into cancer cells, it can still indirectly contribute to tumor progression through paracrine crosstalk. We believe that our data provides evidence that encourages future studies to investigate whether antibiotic-mediated therapy regimes in conjunction with chemotherapy or inhibition of cytokine and chemokine receptors may prove beneficial for treating these as well as other cancers that contain a tumor microbiome which may act to exacerbate disease.

## Materials & Methods

### Culturing Bacteria

*Fusobacterium nucleatum subsp. nucleatum* ATCC 23726 (referred to as *Fnn*) and its Fap2 surface adhesin deletion mutant, *F. nucleatum Δfap2* (referred to as *Fnn Δfap2*), were grown as detailed previously^37^. Bacterial colonies are initiated on solid agar plates made with Columbia Broth (Gibco) substituted with hemin (5 μg/mL) and menadione (0.5 μg/mL) (CBHK) and grown in anaerobic conditions (90% N_2_, 5% H_2_, 5% CO_2_) at 37°C for two days. Single colonies were then retrieved and added to CBHK media to initiate the liquid culture and were grown for ~16 hours at 37°C anaerobically. The optical density (OD) at 600 nm was then measured (~0.6-0.8) to obtain bacteria at the mid-exponential growth phase of the culture. Bacterial counts were obtained based on standardized curves created for the spectrophotometer. All experiments were carried out with a 50:1 multiplicity of infection (MOI) which is the bacteria to host cell ratio.

To stain the bacteria, 1 mL of the bacterial cell suspension was collected and spun down at 1500g for 3 minutes. The pellet was resuspended in 100 μL CBHK media and then stained with FM 1-43FX lipophilic styryl dye (Invitrogen F35355) (5 μg/mL) for 5 minutes. This stains the outer membrane of the bacteria to emit green fluorescence upon excitation. The stained cells were subsequently spun down at 1500g for 3 minutes, washed with 500 μL media, and again resuspended in 1 mL of media and used for experiments.

*Escherichia coli* TOP10 was grown in the shaker incubator at 37°C in Luria Broth (LB) overnight. Bacterial culture was then accordingly diluted to OD_600nm_ of 0.6-0.8 and used for experiments.

### Culturing Normal and Cancerous Epithelial Cells

Cancerous epithelial cells were purchased from ATCC and grown on tissue culture-treated plates and flasks in their respective cell culture media. BxPC3 (ATCC CRL-1687) cells were grown in RPMI-1640 supplemented with 10% FBS, and Panc1 (ATCC CRL-1469) cells were grown in DMEM supplemented with 10% FBS. Normal primary pancreatic epithelial cells were purchased from CellBiologics (H-6037), and grown in epithelial cell culture medium supplemented with 5% FBS (Cell Biologics H-6621). Additional cell lines HPAC (ATCC CRL-2119) and Capan1 (ATCC HTB-79) cells were grown in DMEM:F12 supplemented with 10% FBS and IMDM supplemented with 20% FBS, respectively. All cell culture media contained 1% penicillin and streptomycin, but infections were performed in serum-free and antibiotic-free media. Cells were grown at 37°C with 5% CO_2_ and were passaged every 2-3 days based on their confluency with gentle trypsinization (0.25% trypsin with EDTA) and reseeding. The cells used for the experiments did not exceed passage 12. For infection experiments, cells were grown to confluency in 6, 12, 24, and 96 well plates.

### Scanning Electron Microscopy

BxPC3 cells were seeded on coverslips and allowed to adhere overnight. The samples were then infected with *Fnn* for 2 hours at 50:1 MOI (multiplicity of infection). Preparation for scanning electron microscopy was performed as previously described^83^. In brief, samples were fixed for 24 h with 4% paraformaldehyde and 1% glutaraldehyde in PBS, post-fixed for 45 min with 1% osmium tetroxide in distilled water, and subsequently dehydrated in serial dilutions of ethanol of 25%, 50%, 70%, 95%, 100%, and 100% before critical point drying the samples. Samples were then sputter-coated with iridium fast coating (5 nm) on a Leica EM ACE600 and imaged on LEO (Zeiss) 1550 Field Emission SEM.

### Flow Cytometry

*Fnn* and *Fnn Δfap2* were obtained and first stained with FM 1-43FX as detailed above to facilitate green fluorescence detection. The epithelial cells were then infected in serum-free and antibiotic-free media with the stained bacteria at 50:1 MOI for 1 hour and 4 hours. The cells were then washed twice with PBS, trypsinized, and collected for flow cytometry and cell sorting. The population of cells containing a mix of non-infected and infected cells was then loaded onto an S3e flow cytometer (Bio-Rad). 50,000 cells were analyzed using single-cell gates to measure the median green fluorescence induced by intracellular *Fnn*. FlowJo10 was then used to determine the median fluorescence of the samples and the data was transferred to GraphPad Prism for statistical analysis.

To visualize intracellular bacterial counts based on fluorescence intensity, the cells were sorted based on gates created within the infected cell population for low, medium, and high fluorescence. The sorted cells were collected in 100% FBS, transferred to growth media, and then plated on coverslips for imaging. After adherence, the cells were stained with MemBrite 568/580 (Biotium BTM30095) and NucBlue (ThermoFisher Scientific R37605) before being fixed with 10% formalin. The cells were imaged on a Zeiss LSM 800 using a 63X oil objective and z-stacks were obtained to recreate the 3D structure of the infected cells on Zen Blue.

### Cell Proliferation Assays

Proliferation assays were performed using XTT Assay Kit (ATCC 30-1011K) and BrdU Cell Proliferation Assay Kit (K306, BioVision) based on the protocols provided. Briefly, epithelial cells were seeded on 96 well plates at optimized cell numbers (10,000 cells/well) and allowed to adhere for 12 hours in their respective cell culture media. The media was then replaced with the treatment (conditioned media or cell culture media with added cytokines 200 pg/mL, each supplemented with 0.5% FBS) or the cells were directly infected on the 96-well plate at 50:1 MOI for 4 hours. The cancer cells were further incubated for 48 hours, and the primary normal cells were incubated for 24 hours at 37°C, which was optimized to account for their doubling time. The recombinant cytokines that were used were Recombinant Human GM-CSF Protein (R&D Systems, 215GM), Recombinant Human CCL20/MIP-3α Protein (R&D Systems, 360-MP), IL-8 Monocyte Recombinant Human Protein (ThermoFisher, PHC0884), and Recombinant Human CXCL1/GROα Protein (R&D Systems, 275-GR).

For the XTT assay, following the 24/48-hour incubation with conditioned/cytokine media, 50μL of activated XTT reagent is added to each well and the resulting color change was monitored for 2-24 hours by measuring the absorbance at 475 nm and 660 nm. For the BrdU assay, 10X BrdU solution is added to the treated cells to a final concentration of 1X and incubated for 4 hours. The medium was then removed and the cells were fixed with the Fixing/Denaturing solution for 30 minutes. Next, BrdU Detection Antibody was added and incubated for 1 hour with shaking, followed by washing thrice with Wash Buffer. Anti-mouse HRP-linked Antibody Solution was added to the wells and it was incubated for 1 hour and washed thrice again with Wash Buffer. Finally, TMB substrate was added to the wells and the color change was monitored for 5-20 minutes and the final absorbance was measured at 450 nm.

### Preparing Conditioned Media

The epithelial cells were grown to confluence in their respective media supplemented with 10% FBS and 1% penicillin-streptomycin in T-75 flasks or 12 well plates. One set of T-75 flasks/wells was then used for infection and another set was used as a control without infection. The flasks were washed with PBS twice and replaced with serum-free and PS-free media. For testing the impact of nPEC conditioned media on cancerous cells, serum-free and PS-free media of the cancer cell line was used. The flasks were then infected with *Fnn* at 50:1 MOI for 4 hours. The media from both uninfected and infected cells was retrieved from the flasks and filtered through a 0.22 μm syringe filter (Millipore Sigma). For proliferation assays, conditioned media was used directly without concentration. For migration assays, the extracted media was concentrated to 16X, as determined from preliminary trials, using a 3000 MWCO Concentrator (Amplicon Millipore Sigma) by spinning the samples at 3000g for ~1.5 hours at 4°C. The samples were immediately used for the Transwell assays and ELISA.

### Cell Migration Assays

Cell migration assays were performed on 8μm Transwells in 24-well plates (Corning CLS3422). Cells were first trypsinized (0.25% Trypsin in EDTA) and stained using CellTracker Red CMPTX (ThermoFisher C34552) for 45 minutes in serum-free media. Matrigel (Corning CB40234A) was diluted to a concentration of 250 μg/mL in serum-free media and 100μL was added to the upper chamber of each Transwell and incubated for 25 minutes at 37°C. The stained cells were resuspended to 2 million cells/mL in media containing 0.5% FBS and 100μL of the cell suspension was added on top of the Matrigel and incubated for 1 hour. The control and chemoattractant solutions were prepared in serum-free media or media with 0.5% FBS (100 ng/mL) and 600μL of the chemoattractant solution was added to the outer chamber of the Transwell. In the case of migration in response to conditioned media, 600μL of concentrated conditioned media was added to the outer chamber of the Transwell. The Transwells were then incubated for the respective time points (12/16/24) hours at 37°C/5% CO_2_. The plate was collected and the cells in the upper chamber were removed using a cotton-tip applicator. The Transwells were transferred to a plate containing 10% formalin and fixed for 20 minutes. This was washed with PBS thrice and stained with DAPI (1:5000) (Sigma Aldrich D9542) in a permeabilization buffer (10 mL PBS + 0.5% Triton-X + 200 mg Bovine Serum Albumin) overnight at 4°C. Transwells were then washed thrice with PBS and stored at 4°C. The Transwells were imaged on a Zeiss LSM 800 confocal microscope using a 10X objective. The cells were counted using a custom-developed ImageJ protocol.

### Cytokine Arrays

Cytokine arrays were obtained using the Proteome Profiler Human XL Cytokine Array Kit (R&D Systems Inc. ARY022B) that screens for 105 cytokines. Briefly, cells were grown to confluence in 6 well plates. The cells were infected with *Fnn, E. coli*, and *Fnn Δfap2* at 50:1 MOI for 4 hours at 37°C. Infection with *E. coli* TOP10 was used as a non-invasive infection control. The samples were first collected and filtered using a Spin-X column (Millipore Sigma) to remove bacteria within the sample. The membranes from the kit were first blocked using the Assay buffer for 1 hour and then incubated with the samples overnight at 4°C on a rocking platform shaker. Each membrane was then washed using the Wash buffer, and incubated with the Detection antibody cocktail for 1 hour. This was followed by washing thrice with Wash Buffer again and incubating with Streptavidin-HRP for 30 minutes. After a final washing procedure, Chemi-Reagent Mix was added to each membrane and it was covered for 1 minute. Finally, the membranes were transferred to an autoradiography film cassette and exposed to X-rays for 1-10 minutes.

### Chemokine Quantitation

ELISA was used to quantify cytokines IL-8, CXCL1, MIP-3α, and GM-CSF. The assays were performed using Duo Kits (R&D Systems DY208, DY275, DY360, and DY215, for the 4 cytokines respectively). Epithelial cells were grown to confluence on 12 well plates and infected with *Fnn, E. coli*, and *Fnn Δfap2* at 50:1 MOI for 4 hours at 37°C. The ELISA plate was first coated with the capture antibody overnight and blocked the next day with the reagent diluent for 1 hour. Standards were prepared and conditioned media obtained from infected cells was filtered using a Spin-X column and then added to the wells, diluted, if necessary, with the reagent diluent. After a 2-hour incubation at room temperature, the plate was washed thrice with Wash buffer, and then incubated with the Detection Antibody for 2 hours at room temperature. The plate was again washed and incubated with Streptavidin-HRP for 30 minutes. Finally, the plate was washed and incubated with the Substrate solution for 20-30 minutes and the reaction was stopped by adding the Stop solution. The final absorbance was measured at 450 nm on a spectrophotometer.

### Chemokine Depletion Assays

Conditioned and concentrated media was obtained as previously described. Human GM-CSF biotinylated antibody (R&D Systems, BAM215) was added to the medium to a final concentration of 90 ng/ml and incubated at room temperature with gentle shaking for 3 hours. Next, magnetic streptavidin particles (Sigma-Aldrich, 11641778001) were first washed twice with PBS and spun down at 1500 g and resuspended in the cell media, and 25 μL/mL was added to the sample media. This was incubated at room temperature for 1 hour. The samples were then spun down at 1500g to pellet the heavier magnetic particles, and the supernatant was collected containing the conditioned medium with reduced cytokines. The medium was used for the proliferation assays and an ELISA was used to quantify and confirm the cytokine depletion.

### Statistical Analysis

Statistical analysis was performed on GraphPad Prism 9 using t-tests and ANOVA followed by Dunnett’s multiple comparisons test, with P-value significance denoted by ns (not significant) for P>0.05, * for P≤0.05, ** for P≤0.01, *** for P≤0.001, and **** for P≤0.0001. All samples for analysis were collected in independent triplicates. Plots were created on GraphPad Prism 9 and figures were designed on Affinity Designer.

## Acknowledgments

This project was supported by the NIH R21 Exploratory/Developmental Research Grant 1R21CA238630-01A1 (Verbridge, Slade), NSF Career Award CBET-1652112 (Verbridge), College of Engineering, and the Center for Engineered Health of the Institute for Critical Technologies and Applied Sciences at Virginia Tech. For SEM imaging experiments, we would also like to acknowledge the Nanoscale Characterization and Fabrication Laboratory, which is supported by the Virginia Tech National Center for Earth and Environmental Nanotechnology Infrastructure (NanoEarth), a member of the National Nanotechnology Coordinated Infrastructure (NNCI), supported by NSF (ECCS 1542100 and ECCS 2025151). The summary figure was designed using Biorender.com (Toronto).

## Author Contributions

Conceptualization: BU, SSV, DJS

Formal Analysis: BU, RA

Funding acquisition: SSV, DJS

Investigation: BU, TTDN, AU, RA, LMR, PS, SJ

Methodology: BU, TTDN, AU, RA, LMR

Project administration: SSV, DJS, JMM

Supervision: SSV, DJS, JMM

Visualization: BU

Writing - original draft: BU

Writing - review and editing: BU, TTDN, AU, RA, LMR, JMM, DJS, SSV

## Materials & Correspondence

Scott S. Verbridge

