## Supplemental Figures for "*Fusobacterium nucleatum* induces pancreatic cancer cell proliferation and migration by regulating host autocrine and paracrine signaling"

### *Supplementary Figures*

#### Supplementary Table 1

| Cell Line | Age | Sex | Derivation | Metastasis | Proliferation | Differentiation | Reference |
|---|---|---|---|---|---|---|---|
| BxPC3 | 61 | Female | Primary Tumor | No | 48-60 h | Moderate to Poor | Tan et al. (1986) PubMed 3754176 |
| Panc1 | 56 | Male | Primary Tumor | Yes | 52 h | Poor | Leiber et al. (1975) PubMed 1140870 |
| Capan-1 | 40 | Male | Liver metastasis | Yes | ND | Well | Kyriazis et al. (1982) PubMed 6278935 |
| HPAC | 64 | Female | Primary Tumor | ND | 41 h | Moderate | Gower et al. (1994) |

**Supplementary Table 1:** Cancer cell lines and their noted features.

#### Cytokine Arrays

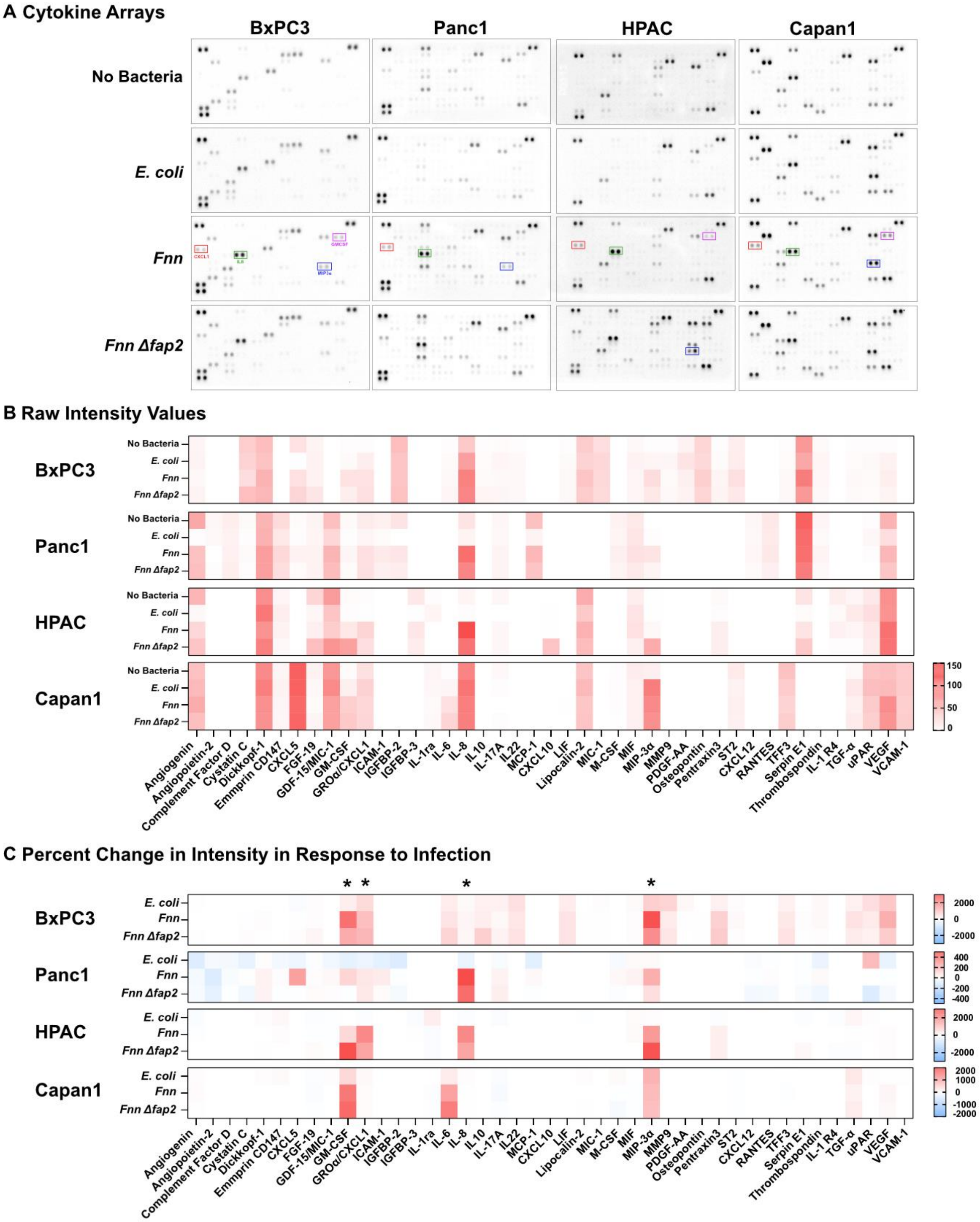


**Supplementary Figure 1: Cytokine arrays of infected host cell secretions. A.** Cytokine secretions were tested for 4 conditions: No Bacteria, *E. coli*, *Fnn, and Fnn ∆fap2*. using the XL Cytokine Array Kit (R&D Systems). **B.** Raw intensity values based on the blots were determined on ImageJ and the intensities are plotted as a heat map. **C.** Percent change in intensity relative to constitutive secretion was determined for all the infection conditions.

#### Cytokine Secretions

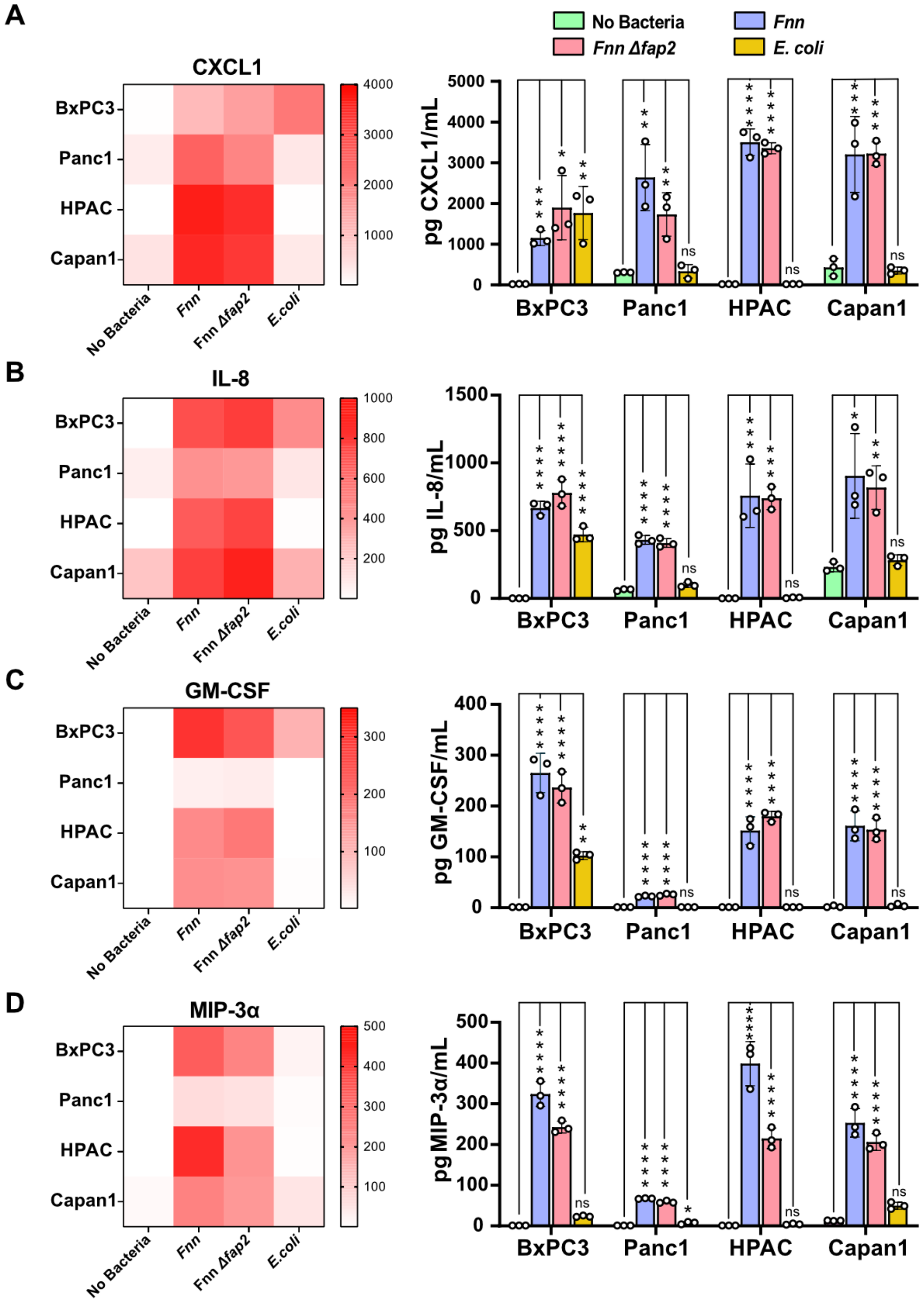


**Supplementary Figure 2:** ELISA was used to quantify the levels of secretion of **A.** CXCL1, **B.** IL-8, **C.** GM-CSF, and **D.** MIP-3α for all 4 pancreatic cancer cell lines tested.

#### Cytokine impact on proliferation

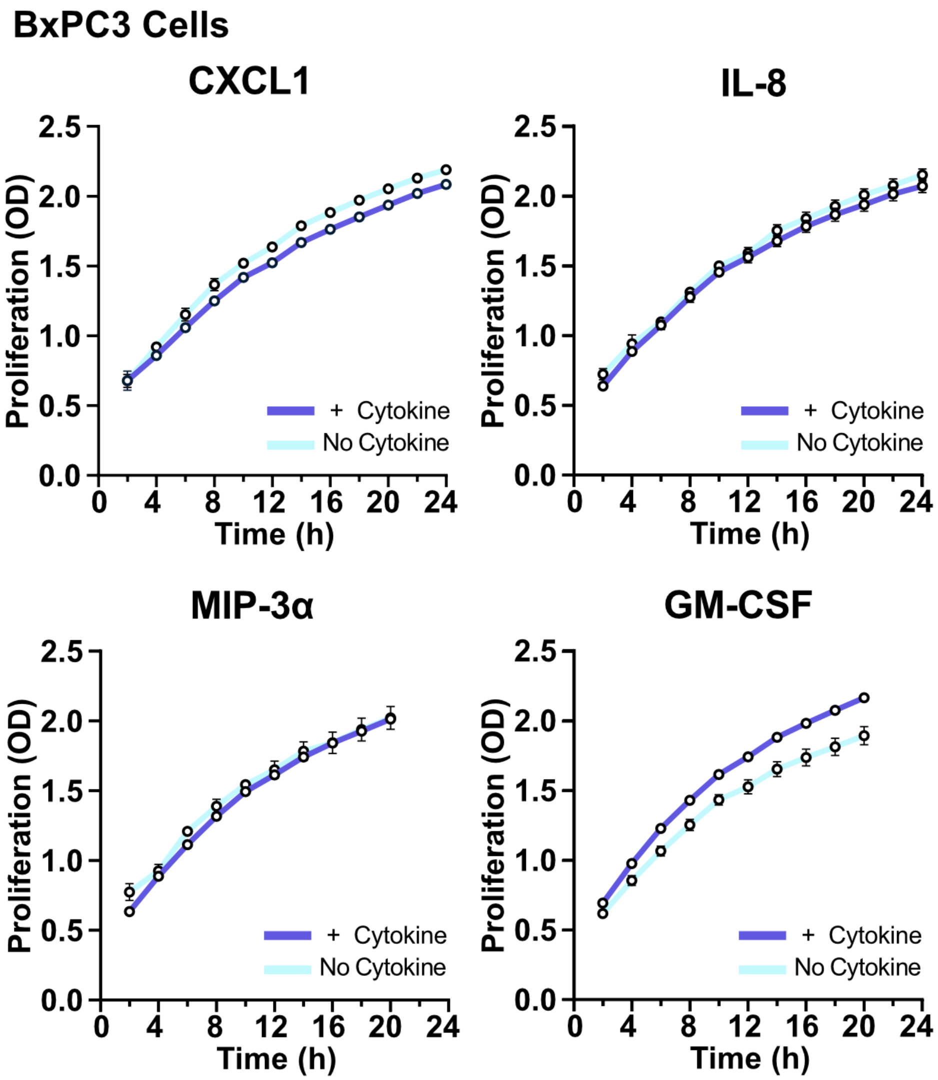


**Supplementary Figure 3: GM-CSF contributes to increased proliferation of BxPC3 cells.** Preliminary studies were conducted on BxPC3 cells to identify the impact of the 4 cytokines individually on proliferation using the XTT assay over 24 hours. GM-CSF at 200 pg/mL showed a significant increase in proliferation compared to the other cytokines.

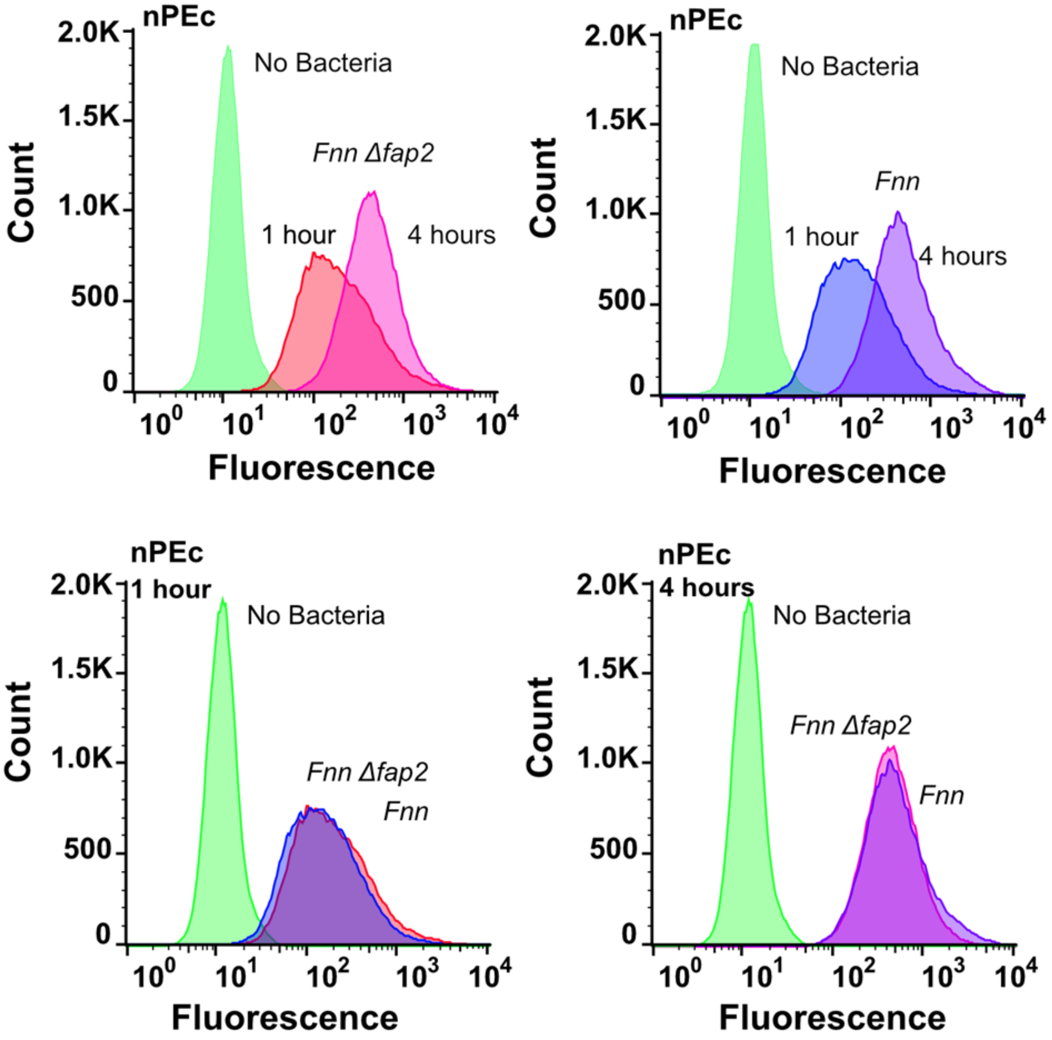


**Supplementary Figure 4: Fap2 does not impact *F. nucleatum* infection of normal pancreatic cells.** Flow cytometry data of normal pancreatic epithelial cells (nPEc) after infection with *Fnn* and *FnnΔfap2* for 1 hour and 4 hours of infection.
